# On the mechanism of bilayer separation by extrusion; or, why your large unilamellar vesicles are not really unilamellar

**DOI:** 10.1101/764274

**Authors:** Haden L. Scott, Allison Skinkle, Elizabeth G. Kelley, M. Neal Waxham, Ilya Levental, Frederick A. Heberle

## Abstract

Extrusion through porous filters is a widely used method for preparing biomimetic model membranes. Of primary importance in this approach is the efficient production of single bilayer (unilamellar) vesicles that eliminate the influence of interlamellar interactions and strictly define the bilayer surface area available to external reagents such as proteins. Sub-microscopic vesicles produced using extrusion are widely assumed to be unilamellar, and large deviations from this assumption would dramatically impact interpretations from many model membrane experiments. Using three probe-free methods—small-angle X-ray and neutron scattering (SAXS and SANS) and cryogenic electron microscopy (cryoEM)—we report unambiguous evidence of extensive multilamellarity in extruded vesicles composed of neutral phosphatidylcholine lipids, including for the common case of neutral lipids dispersed in physiological buffer and extruded through 100 nm diameter pores. In such preparations, only ~35% of lipids are externally accessible, and this fraction is highly dependent on preparation conditions. Charged lipids promote unilamellarity, as does decreasing solvent ionic strength, indicating the importance of electrostatic interactions in determining the lamellarity of extruded vesicles. Smaller extrusion pore sizes also robustly increase the fraction of unilamellar vesicles, suggesting a role for membrane bending. Taken together, these observations suggest a mechanistic model for extrusion, wherein formation of unilamellar vesicles involves competition between bilayer bending and adhesion energies. The findings presented here have wide-ranging implications for the design and interpretation of model membrane studies, especially ensemble-averaged observations relying on the assumption of unilamellarity.

**STATEMENT OF SIGNIFICANCE:** Extruded vesicles are a ubiquitous tool in membrane research. It is widely presumed that extrusion produces unilamellar (i.e., single bilayer) vesicles, an assumption that is often crucial for data analysis and interpretation. Using X-ray and neutron scattering and cryogenic electron microscopy, we show that a substantial amount of lipid remains inaccessible after extrusion due to an abundance of multilamellar vesicles (MLVs). While this is a general phenomenon for neutral lipids, MLV contamination can be reduced by several complementary approaches such as including charged lipids in the mixture, reducing the ionic strength of the aqueous medium, and reducing the extrusion pore size. These observations together suggest a mechanism by which extrusion strips MLVs of their layers.

---

Biomimetic model membranes enable carefully controlled, systematic experiments that can inform on complex functionality in living membranes. Many insights into biological membrane structure (1,2), shape transformations (3), dynamics (4), and phase behavior (5–7) derive from studies of chemically defined synthetic bilayers. These systems have also contributed greatly to our understanding of the interactions between proteins and biomembranes (8).

While synthetic membranes can be produced in a variety of sizes and geometries, perhaps the most widely used experimental system consists of extruded vesicles (EVs) with diameters on the order of 100 nm. These vesicles are smaller than the resolution limit of light microscopy, but their size distributions can be measured with dynamic light scattering (DLS). Because of their uniform size and physical properties, extruded vesicles are readily amenable to ensemble-averaged spectroscopic and scattering techniques, as well as a vast array of biochemical assays. For many experiments, a convenient assumption is that half of the total lipid is exposed to the external solvent where it can interact with reagents in the extravesicular space. Unfortunately, few techniques are directly sensitive to the presence of inaccessible lipid layers in vesicle samples. Instead, uniformity in DLS-reported vesicle size distributions is often taken as evidence of successful extrusion and consequently, of a sufficiently unilamellar sample; indeed, such preparations are typically referred to as Large Unilamellar Vesicles, or LUVs. In this Letter, we show that the assumption of unilamellarity is incorrect for many commonly used vesicle formulations.

Figure 1A shows SAXS scattering intensity versus momentum transfer *q* for POPC dispersions in PBS buffer at 25°C (all membranes in this study are measured in the fluid lamellar phase). Without extrusion, a characteristic set of broad, equally spaced peaks (in this case, at *q* = 0.10 Å^−1^, 0.20 Å^−1^, and 0.30 Å^−1^) emerges from lobes of diffuse scattering. These peaks are the first three orders of Bragg reflections from a repeating structure in the sample with a spacing of *d* = 2π/*q*_1_ = 62.6 Å. Given the well-known tendency of lipid films to swell into multilamellar vesicles (MLVs) consisting of multiple concentric bilayers separated by water, the observation of Bragg peaks corresponding to the lamellar repeat distance (i.e., one bilayer plus an interstitial water layer) is expected. The surprising result is that standard extrusion protocols believed to produce 100 nm unilamellar vesicles do not eliminate these reflections (Fig. 1A). Though the peaks in the EV sample are broader and less intense, their prominence indicates that substantial multilamellar structures remain even after extrusion.

**Figure 1.**
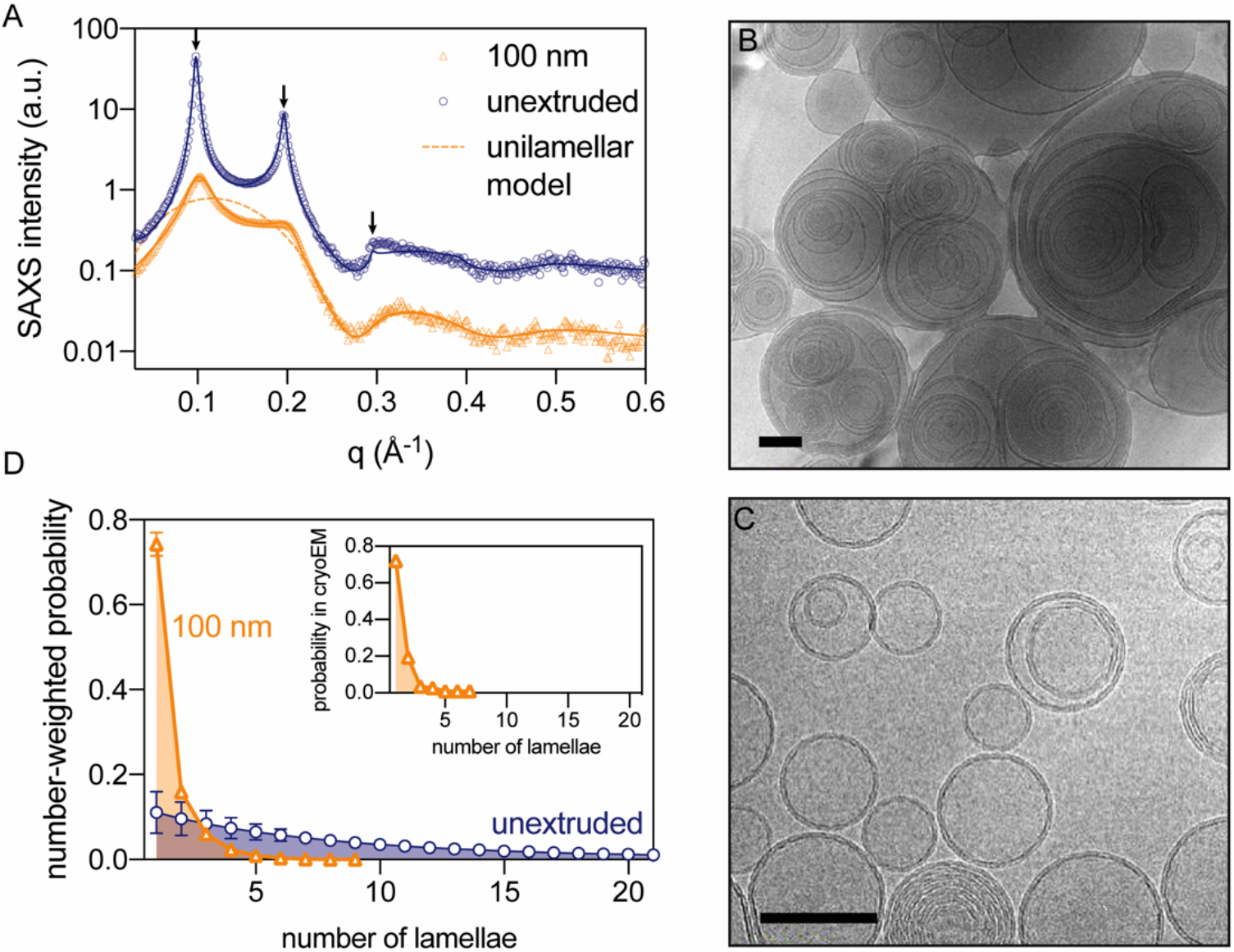
SAXS and cryoEM of lipid vesicles reveal extensive multilamellarity after extrusion. (A) SAXS intensity vs. *q* for unextruded POPC suspensions (blue circles) and POPC hydrated in PBS buffer and extruded through 100 nm pores (orange diamonds). The solid curves are fits to a scattering model that accounts for bilayer structure and interlamellar correlations, which appear as regularly spaced peaks (arrows). The dashed curve shows the expected form factor of a purely unilamellar vesicle. (B) cryoEM images of unextruded and (C) 100-nm pore extruded POPC suspensions (scale bar 100 nm). (D) Number-weighted distributions of vesicle lamellarity obtained from model fitting to SAXS data. (inset) Probability distribution of extruded vesicle lamellarity for similar samples obtained from cryoEM.

We confirmed the presence of MLVs in both non-extruded (Fig 1B) and extruded (Fig 1C) samples by direct, real-space cryoEM imaging. Multilamellar and unilamellar vesicles in the extruded sample are comparable in size (Fig. 1C), suggesting that all vesicles passed through the extrusion pores (as opposed to leaking through the filter edges), and emphasizing that a uniform size distribution in DLS measurements does not imply the absence of MLVs in extruded samples. Indeed, we find that the extent of lamellarity (determined from SAXS data as described below) is only weakly correlated with average vesicle size, and is not correlated with the size polydispersity, both determined from DLS (Table S2 and Fig. S1).

The combination of Bragg peaks in scattering data and direct observation in cryoEM images provides unambiguous proof of multilamellar structures in EV preparations, an outcome that is usually undesirable and often overlooked. To quantify the extent of multilamellarity, scattering data were fit to a model combining a single lipid bilayer form factor and a multilamellar structure factor parameterized with a modified exponential distribution for the number of lamellae (see Supporting Materials and Methods, Section S1 for details of the model and fit procedure). The solid lines in Fig. 1A demonstrate that the model successfully captures both the diffuse lobes of scattering originating from the bilayer structure, as well as the broad reflections generated by the lattice of stacked lamellae. The dashed line shows the expected scattering from purely unilamellar vesicles. Figure 1D compares distributions of the number of lamellae obtained from SAXS analysis to those obtained by counting ~100 vesicles in cryoEM images, revealing excellent agreement between these approaches. As expected, unextruded vesicles showed a wide distribution that narrows and shifts toward fewer lamellae upon extrusion; however, only half of the lipid mass in the extruded sample (53 ± 4%) is in unilamellar structures, as determined from four replicate samples of POPC in PBS buffer (Table S2).

Lamellar distributions enable a straightforward estimate of the fraction of externally accessible lipid *f*_*acc*_ (see Supporting Materials and Methods, Section S1 for details). This parameter is crucial for many experiments, as it reports directly on the fraction of total lipid accessible to externally added reagents such as proteins, quenchers, or small molecules. A perfectly unilamellar vesicle sample would have approximately half of its total lipid exposed on the vesicle surface (*f*_*acc*_ ≈ 0.5). Figure 2 shows that common preparations (e.g., 100 nm vesicles of PC lipid hydrated in physiological buffer) yield *f*_*acc*_ = 0.35, ~30% lower than expected for a unilamellar sample. Put another way, approximately one-third of the lipids expected to be accessible to external reagents are instead entrained in vesicle lumens.

**Figure 2.**
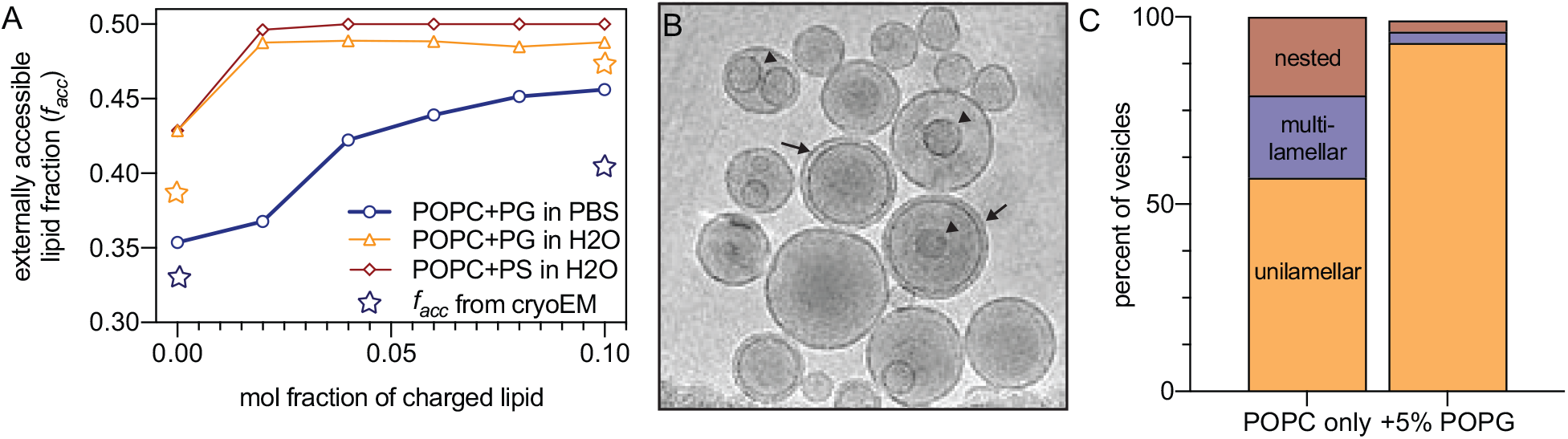
Unilamellarity increases with charged lipids and/or decreasing solvent ionic strength. (A) The fraction of externally accessible lipid *f*_*acc*_ (related to extent of unilamellarity) increases with increasing abundance of charged lipid dopant (POPG or POPS) in POPC bilayers extruded through 100 nm pores. Samples hydrated in PBS (blue circles) show a lower *f*_*acc*_ at all concentrations of charged lipid compared to hydration in pure water. Blue and orange stars represent *f*_*acc*_ calculated from cryoEM images of POPC in PBS and POPC+PG in H_2_O, respectively. (B) Typical cryoEM image of POPC vesicles extruded (100 nm) in water (scale bar 100 nm). Arrows highlight multilamellar structures, while arrowheads highlight nested vesicles. (C) Distribution of structures in 100 nm vesicles extruded in water obtained from cryoEM.

We observed qualitative agreement between *f*_*acc*_ calculated from SAXS and cryoEM images (stars in Fig. 2). Images also revealed unexpected complexity, including bilamellar structures consisting of larger vesicles enveloping smaller “nested” vesicles (Fig 2B, arrowhead), which comprised ~20% of 100 nm POPC vesicles (Fig. 2C). Scattering analyses likely do not fully capture the abundance of these structures, which manifest in *I*(*q*) as a gentle oscillation in the low-*q* regime where the intensity is also influenced by vesicle concentration and size distribution, making it difficult to unambiguously assign scattering features. These nested structures likely explain why *f*_*acc*_ reported by cryoEM was systematically lower than estimates from SAXS (Fig. 2A).

It has previously been reported that a small amount of charged lipid can reduce or eliminate Bragg reflections in scattering data (9,10). Hypothesizing that interlamellar repulsion would promote unilamellarity, we investigated the influence of charged lipids and buffer ionic strength on *f*_*acc*_. Inclusion of negatively charged POPG in POPC bilayers at concentrations up to 10 mol% decreased multilamellarity and increased lipid exposure, as evidenced by both SAXS and cryoEM (Fig 2A&C, Table S2, Fig. S2), with a similar effect observed in SANS data from DMPC/DMPG bilayers (Table S3, Fig. S3B). Greater unilamellarity (i.e., higher *f*_*acc*_) was also achieved without changing membrane lipid composition, but instead by reducing solvent ionic strength, as shown in Fig. 2 for POPC hydrated with water. The absence of screening counterions greatly increased *f*_*acc*_ even in vesicles composed of zwitterionic lipids with no net charge, and further inclusion of as little as 2 mol% POPG or POPS resulted in essentially purely unilamellar vesicles. In good agreement with SAXS, 93% of the POPG-containing vesicles extruded in water were unilamellar in cryoEM images (Fig 2C). Importantly, neither adding charged lipid at < 10 mol% nor changing buffer composition caused significant changes to the area per lipid (Table S2), suggesting that overall bilayer structure was unaffected. Thus, while near-complete unilamellarity with minimal bilayer perturbation can be achieved in samples containing charged lipid and hydrated in pure water, the absence of either factor yields a substantial amount of inaccessible lipid even after extrusion through 100 nm pores. Together, the effects of charged lipid and ionic strength point to a role for electrostatic repulsion in the production of unilamellar vesicles by extrusion.

We next investigated approaches to increase unilamellarity without altering the chemical composition of the bilayer or its aqueous solvent. Figure 3 shows the influence of extrusion pore size for POPC bilayers with or without 5 mol% POPG (see also Table S2 and Fig. S4). In both cases, decreasing the pore size increased the fraction of externally accessible lipid, with similar effects observed by SANS for DLPC and DMPC (Table S3, Fig. S3A, Fig. S5). The largest extrusion pore sizes (400 nm) produced vesicles that were difficult to categorize by cryoEM, containing complex multilamellar and nested structures (Fig. 3B, left). In contrast, the smallest pore filters (30 or 50 nm) or sonication produced essentially unilamellar vesicles even in the absence of charged lipids, as evidenced by cryoEM (Fig. 3B), SAXS (Fig. 3A, Table S2), and SANS (Table S3). Taken together, these observations suggest an interplay between interlamellar repulsion and membrane curvature as key determinants of the efficiency of unilamellar extrusion.

**Figure 3.**
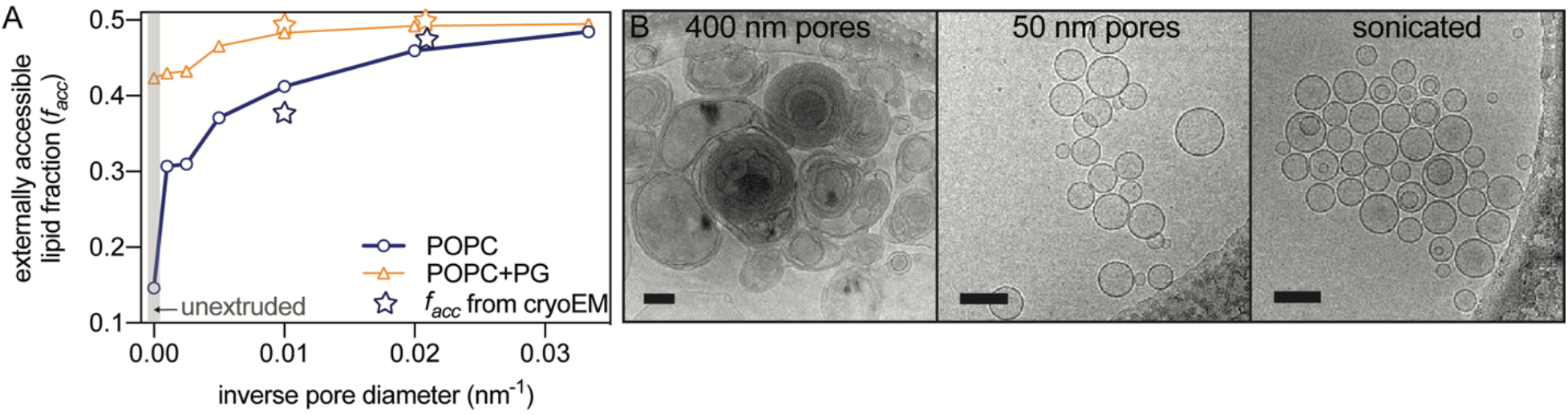
Unilamellarity increases with decreasing extrusion pore size. The fraction of externally accessible lipid increases with decreasing extrusion pore size for POPC vesicles without (blue circles) or with (orange triangles) 5 mol% POPG. Stars represent *f*_*acc*_ calculated from cryoEM images. (B) Typical cryoEM images of POPC vesicles extruded in water through 400- and 50-nm pore filters, as well as after sonication (scale bar 100 nm).

Finally, we explored the influence of lipid chain structure on lamellarity. For eight different phosphatidylcholines including saturated, unsaturated, and mixed chain species, *f*_*acc*_ ranged from 0.39 to 0.46 after extrusion through 100 nm pores (Table S2, Fig. S6B). Surprisingly, egg sphingomyelin (ESM), also a neutral lipid, was significantly more unilamellar than any of the PC lipids (*f*_*acc*_ = 0.49). To explore the possible mechanisms underlying these results, we evaluated correlations between *f*_*acc*_ and membrane physical properties, including total number of carbons in the lipid chains, total number of double bonds in the chains, area per lipid, hydrocarbon thickness, melting temperature, and bending rigidity (Table S5, Fig. S7). The strongest correlation was with the bending energy: *f*_*acc*_ exhibits an approximately linear dependence on the bending modulus *k*_*c*_ determined from molecular simulations (11).

Taken together, the above results suggest a mechanism in which the lamellarity of an extruded sample is influenced by a competition between bilayer bending and adhesion energies. During extrusion, stacked bilayers must collectively bend to enter the cylindrical extrusion pore. Assuming mechanically uncoupled bilayers, the energy required to bend a multilamellar stack scales with both the number of bilayers and the membrane’s bending rigidity and is inversely related to the radius of curvature. Rather than bending, one or more of the bilayers can instead separate from the stack and enter the pore with a smaller bending penalty, a process that is opposed by the interbilayer adhesion energy. Any factor that increases the cost of bending—for example, a larger intrinsic bending stiffness or a smaller pore radius—also increases the likelihood of lamellar separation. Similarly, any factor that lowers the bilayer adhesion energy, such as increasing interlamellar repulsion with charged lipids (12), also favors separation of layers and should therefore increase the fraction of unilamellar vesicles after extrusion. It is likely that bending energy also contributes to an increased probability of unilamellar vesicles upon hydration (i.e., prior to extrusion), as recently proposed (13). Indeed, we found that the lipid with the largest bending modulus (ESM) had the greatest fraction of unilamellar vesicles both before and after extrusion (Table S2). However, sphingomyelin appears to be an outlier in this regard; for the PC lipids we examined, there was no correlation between *f*_*acc*_ before and after extrusion (Table S2, Fig. S6), suggesting that their final lamellar distribution depends predominantly on events occurring during extrusion.

The predictive power of the proposed mechanism relies on the ability to independently control interlamellar forces and bending rigidity, a requirement that is likely not strictly met in any real experiment. In some cases, the different influences reinforce each other: for example, it is well established that adding charged lipid to neutral bilayers increases lamellar separation which in turn reduces bilayer adhesion (14), and there is evidence that charged lipid can also increase membrane bending rigidity (15, 16). Both of these factors favor unilamellarity, consistent with experimental observations reported here. On the other hand, the effect of salts may be difficult to predict due to their complicated (and in many cases, uncertain) influence on interbilayer forces and membrane mechanical properties. For example, the addition of monovalent salt has been shown to reduce attractive van der Waals forces in neutral bilayers (17,18), but also decrease interbilayer separation (and thereby increase adhesion) in charged bilayers through screening effects (the Debye length decreases by three orders of magnitude between pure water and physiological salt concentration). Moreover, NaCl has been shown to increase the bending rigidity of POPC multilayers (19) but decrease the bending rigidity of POPC giant unilamellar vesicles (16). Divalent cations are even more complicated due to their ability to tightly bind acidic headgroups (20). In short, the proposed mechanism should be used with appropriate caution; we recommend an approach of experimentally validating the effect of any sample additive using the methods described here.

The presence of multilamellar structures in extruded vesicles has been sporadically reported in the literature, often evidenced by their clear experimental signature in scattering studies (9,13,21). Some factors that can potentially influence multilamellarity are the number of freeze-thaw cycles (22), membrane lipid composition (9), and modification of lipid headgroups, for example by PEG polymers (13,22–24). This work expands on these observations in several important ways. First, we present a general scattering model that enables robust quantification of the accessible lipid fraction from SANS or SAXS data, and we validate the model-dependent analysis with direct cryoEM imaging, a real-space technique that provides a particularly straightforward means to assess lamellarity. Notably, imaging revealed a nested vesicle structure difficult to detect with scattering that seems to be a preferred outcome for some samples; the mechanism responsible for creating this structure is not immediately obvious and warrants further investigation. Second, by examining nine different neutral lipids, we find that while residual MLVs are generally present after extrusion, their abundance differs in a manner that likely depends on membrane mechanical properties and possibly other factors. Finally, we identify three methods for increasing the fraction of unilamellar vesicles: (a) adding small amounts of charged lipid; (b) decreasing solution ionic strength; and (c) reducing extrusion pore size. Practically, these findings highlight a major potential artifact in very commonly used experimental preparations and provide a complementary set of strategies for mitigating the problem of MLV contamination in extruded samples. In addition, they point toward a mechanism by which extrusion strips MLVs of their layers (or in many cases, fails to do so) that can be further explored with theory, simulation and experiment.

## SUPPORTING MATERIAL

Supporting Materials and Methods, five tables, and seven figures are available at http://www.biophysj.org/biophysj/supplemental/XXXX.

## AUTHOR CONTRIBUTIONS

F.A.H., I.L., E.G.K., and M.N.W. designed the research. H.L.S., A.S., E.G.K., M.N.W., and F.A.H. conducted experiments. H.L.S., A.S., E.G.K., M.N.W., I.L., and F.A.H. analyzed data. F.A.H. and I.L. wrote the article. H.L.S., A.S., E.G.K., M.N.W., I.L., and F.A.H. revised the manuscript.

## ACKNOWLEDGMENTS

FAH and HLS were supported by National Science Foundation (NSF) Grant No. MCB-1817929. IL and AS were supported by the National Institutes of Health (NIH)/National Institute of General Medical Sciences (GM114282, GM124072, GM120351), the Volkswagen Foundation (grant 93091), and the Human Frontiers Science Program (RGP0059/2019). MNW was supported by an endowment from the William Wheless III Professorship and acknowledges NIH Shared equipment grants for the Polara (S10 RR16714) and K2 Summit (S10 OD016279) with additional support provided through the Structural Biology Center at The University of Texas Medical School at Houston. Access to the NGB30 SANS Instrument was provided by the Center for High Resolution Neutron Scattering, a partnership between the National Institute of Standards and Technology and the NSF under agreement No. DMR-1508249. SAXS measurements were supported by Department of Energy scientific user facilities at Oak Ridge National Laboratory (ORNL), and DLS measurements were supported by the Biophysical Characterization Suite of the Shull Wollan Center at ORNL.

## SUPPORTING CITATIONS

References (25-35) appear in the Supporting Material.

## REFERENCES

1. Nagle, J. F., and S. Tristram-Nagle. 2000. Structure of lipid bilayers. Biochim. Biophys. Acta. 1469:159–195.

2. Heberle, F. A., Pan, J., Standaert, R. F., Drazba, P., Kučerka, N., and J. Katsaras. 2012. Model-based approaches for the determination of lipid bilayer structure from small-angle neutron and X-ray scattering data. Eur. Biophys. J. 41:875–890.

3. McMahon, H. T., and E. Boucrot. 2015. Membrane curvature at a glance. J. Cell Sci. 128:1065–1070.

4. Rawicz, W., Olbrich, K. C., McIntosh, T., Needham, D., and E. Evans. 2000. Effect of chain length and unsaturation on elasticity of lipid bilayers. Biophys. J. 79:328–339.

5. G. W. Feigenson. 2009. Phase diagrams and lipid domains in multicomponent lipid bilayer mixtures. Biochim. Biophys. Acta 1788:47–52.

6. D. Marsh. 2009. Cholesterol-induced fluid membrane domains: a compendium of lipid-raft ternary phase diagrams. Biochim. Biophys. Acta 1788:2114–2123.

7. D. Marsh. 2010. Liquid-ordered phases induced by cholesterol: a compendium of binary phase diagrams. Biochim. Biophys. Acta 1798:688–699.

8. Andersen, O. S., and R. E. Koeppe, 2nd. 2007. Bilayer thickness and membrane protein function: an energetic perspective. Annu. Rev. Biophys. Biomol. Struct. 36:107–130.

9. Kučerka, N., Pencer, J., Sachs, J. N., Nagle, J. F., and J. Katsaras. 2007. Curvature effect on the structure of phospholipid bilayers. Langmuir 23:1292–1299.

10. Heberle, F. A., Marquardt, D., Doktorova, M., Geier, B., Standaert, R. F., Heftberger, P., Kollmitzer, B., Nickels, J. D., Dick, R. A., Feigenson, G. W., Katsaras, J., London, E., and G. Pabst. Subnanometer Structure of an Asymmetric Model Membrane: Interleaflet Coupling Influences Domain Properties. Langmuir 32:5195–5200.

11. Doktorova, M., Harries, D., and G. Khelashvili. 2017. Determination of bending rigidity and tilt modulus of lipid membranes from real-space fluctuation analysis of molecular dynamics simulations. Phys. Chem. Chem. Phys. 19:16806–16818.

12. Rand, R. P., and V. A. Parsegian. Hydration forces between phospholipid bilayers. Biochim. Biophys. Acta 988:351–376.

13. Nele, V., Holme, M. N., Kauscher, U., Thomas, M. R., Doutch, J. J., and M. M. Stevens. 2019. Effect of Formulation Method, Lipid Composition, and PEGylation on Vesicle Lamellarity: A Small-Angle Neutron Scattering Study. Langmuir 35:6064–6074.

14. Loosley-Millman, M. E., Rand, R. P., and V. A. Parsegian. 1982. Effects of monovalent ion binding and screening on measured electrostatic forces between charged phospholipid bilayers. Biophys. J. 40:221–232.

15. Mertins, O., and R. Dimova. 2013. Insights on the interactions of chitosan with phospholipid vesicles. Part II: membrane stiffening and pore formation. Langmuir 29:14552–14559.

16. Faizi, H. A., Frey, S. L., Steinkuhler, J., Dimova, R., and P. M. Vlahovska. 2019. Bending rigidity of charged lipid bilayer membranes. Soft Matter 15:6006–6013.

17. J. Marra. 1986. Direct Measurements of Attractive Van der Waals and Adhesion Forces between Uncharged Lipid Bilayers in Aqueous Solutions. J. Colloid Interf. Sci. 109:11–20.

18. Petrache, H. I., Zemb, T., Belloni, L., and V. A. Parsegian. 2006. Salt screening and specific ion adsorption determine neutral-lipid membrane interactions. Proc. Nat. Acad. Sci. USA 103:7982–7987.

19. Pabst, G., Hodzic, A., Strancar, J., Danner, S., Rappolt, M., and P. Laggner. 2007. Rigidification of Neutral Lipid Bilayers in the Presence of Salts. Biophys. J. 93:2688–2696.

20. Rand, R. P., and V. A. Parsegian. 1984. Physical force considerations in model and biological membranes. Can. J. Biochem. Cell Biol. 62:752–759.

21. Pabst, G., Koschuch, R., Pozo-Navas, B., Rappolt, M., Lohner, K., and P. Laggner. 2003. Structural analysis of weakly ordered membrane stacks. J. Appl. Crystallogr. 36:1378–1388.

22. Kaasgaard, T., Mouritsen, O. G., and L. Jorgensen. 2003. Freeze/thaw effects on lipid bilayer vesicles investigated by differential scanning calorimetry. Biochim. Biophys. Acta 1615:77–83.

23. Kenworthy, A. K., Simon, S. A., and T. J. McIntosh. 1995. Structure and phase behavior of lipid suspensions containing phospholipids with covalently attached poly(ethylene glycol). Biophys. J. 68:1903–1920.

24. Hristova, K., and D. Needham. 1995. Phase behavior of a lipid/polymer-lipid mixture in aqueous medium. Macromolecules 28: 99–1002.

25. S.R. Kline. 2006. Reduction and analysis of SANS and USANS data using IGOR Pro. J. Appl. Crystallogr. 39:895–900.

26. Doktorova, M., Heberle, F. A., Marquardt, D., Rusinova, R., Sanford, R. L., Peyear, T. A., Katsaras, J., Feigenson, G. W., Weinstein, H., and O. S. Andersen. 2019. Gramicidin Increases Lipid Flip-Flop in Symmetric and Asymmetric Lipid Vesicles. Biophys. J. 116:860–873.

27. Pencer, J., Krueger, S., Adams, C. P., and J. Katsaras. 2006. Method of separated form factors for polydisperse vesicles. J. Appl. Crystallogr. 39:293–303.

28. De Gennes, P. G., and J. Prost. 1993. The Physics of Liquid Crystals. 2nd ed. Oxford University Press, Oxford, UK.

29. Pabst, G., Rappolt, M., Amenitsch, H., and P. Laggner. 2000. Structural information from multilamellar liposomes at full hydration: full q-range fitting with high quality x-ray data. Phys. Rev. E 62:4000–4009.

30. D.N. Mastronarde. 2005. Automated electron microscope tomography using robust prediction of specimen movements. J. Struct. Biol. 152:36–51.

31. Zheng, S. Q., Palovcak, E., Armache, J. P., Verba, K. A., Cheng, Y., and D. A. Agard. 2017. MotionCor2: anisotropic correction of beam-induced motion for improved cryo-electron microscopy. Nat. Methods 14:331–332.

32. Kučerka, N., Nieh, M. P., and J. Katsaras. 2011. Fluid phase lipid areas and bilayer thicknesses of commonly used phosphatidylcholines as a function of temperature. Biochim. Biophys. Acta 1808:2761–2771.

33. Kučerka, N., Gallova, J., Uhrikova, D., Balgavy, P., Bulacu, M., Marrink, S. J., and J. Katsaras. 2009. Areas of monounsaturated diacylphosphatidylcholines. Biophys. J. 97:1926–1932.

34. Pan, J., Heberle, F. A., Tristram-Nagle, S., Szymanski, M., Koepfinger, M., Katsaras, J., and N. Kučerka. 2012. Molecular structures of fluid phase phosphatidylglycerol bilayers as determined by small angle neutron and X-ray scattering. Biochim. Biophys. Acta 1818:2135–48.

35. Pan, J., Cheng, X., Monticelli, L., Heberle, F. A., Kučerka, N., Tieleman, D. P., and J. Katsaras. 2014. The molecular structure of a phosphatidylserine bilayer determined by scattering and molecular dynamics simulations. Soft Matter 10:3716–3725.

