## Supplementary Information for "On the mechanism of bilayer separation by extrusion; or, why your large unilamellar vesicles are not really unilamellar"

The Supporting Material for

consists of Materials and Methods, 5 tables, and 7 figures.

### S1. MATERIALS and METHODS

*Materials.* Phospholipids 1-palmitoyl-2-oleoyl-sn-glycero-3-phosphocholine (POPC), 1-palmitoyl-2-oleoyl-sn-glycero-3-phospho-(1'-rac-glycerol), sodium salt (POPG), 1-palmitoyl-2-oleoyl-sn-glycero-3-phospho-L-serine, sodium salt (POPS), 1-stearoyl-2-oleoyl-sn-glycero-3-phosphocholine (SOPC), 1,2-dioleoyl-sn-glycero-3-phosphocholine (DOPC), 1,2-dieicosenoyl-sn-glycero-3-phosphocholine (diC20:1-PC), 1,2-dierucoyl-sn-glycero-3-phosphocholine (diC22:1-PC), 1,2-dilauroyl-sn-glycero-3-phosphocholine (DLPC), 1,2-dimyristoyl-sn-glycero-3-phosphocholine (DMPC), 1,1-dimyristoyl-sn-glycero-3-phospho-(1'-rac-glycerol), sodium salt (DMPG), 1,2-dipalmitoyl-sn-glycero-3-phosphocholine (DPPC), and chicken egg sphingomyelin were purchased from Avanti Polar Lipids (Alabaster, AL) as dry powders and used as supplied. The phospholipids were dissolved in HPLC-grade chloroform and stored at −80 °C until use. PBS buffer tablets were from AMRESCO (Solon, OH). Ultrapure $H_2O$ was obtained from a High-Q (Wilmette, IL) or Milli-Q Millipore purification system (Burlington, MA). $D_2O$ (99.9%) was from Cambridge Isotope Laboratories (Tewksbury, MA).

*Sample preparation.* Aqueous lipid dispersions at 20 mg/mL were prepared by first mixing appropriate volumes of lipid stocks in organic solvent with a glass Hamilton syringe. The solvent was evaporated with an inert gas stream followed by high vacuum overnight. The dry lipid film was hydrated above the lipid's main transition temperature for at least 1 h with intermittent vortex mixing. The resulting multilamellar vesicle (MLV) suspension was subjected to at least 5 freeze/thaw cycles using a −80 °C freezer, then extruded through a polycarbonate filter using a handheld mini-extruder (Avanti Polar Lipids) by passing the suspension through the filter 31 times. Unsaturated lipids were extruded at room temperature, and saturated lipids and sphingomyelin were extruded at ~10 °C above their main transition temperature. Because different measurement techniques have different optimal concentration ranges, samples were diluted with water or buffer prior to measurement as described below. Samples were typically measured within a few days of preparation.

*Dynamic Light Scattering (DLS).* Samples were diluted with filtered water to ~ 0.1 mg/mL for DLS measurement in a Brookhaven BI-200SM system equipped with a HeNe laser (Brookhaven Instruments, Holtsville, NY). Typical instrument settings were: goniometer angle 90°; laser power 30 mW; aperture 200 μm; total measurement time 4 min. The vesicle size and polydispersity index reported in Table S2 were obtained from a cumulant analysis performed automatically by the instrument software.

*Small-angle X-ray Scattering (SAXS).* Vesicle samples for small-angle X-ray scattering (SAXS) measurements were prepared at 20 mg/mL as described above and measured using a Rigaku BioSAXS-2000 home source system (Rigaku Americas, The Woodlands, TX) equipped with a HF007 copper rotating anode, a Pilatus 100K 2D detector, and an automatic sample changer. SAXS data were collected at a fixed sample-to-detector distance (SDD) using a silver behenate calibration standard, with a typical data collection time of 30 min for MLVs and 3 h for LUVs. The one-dimensional scattering intensity $I(q)$ [$q = 4\pi \sin(\theta)/\lambda$, where $\lambda$ is the X-ray wavelength and $2\theta$ is the scattering angle relative to the incident beam] was obtained by radial averaging of the corrected 2D image data, an operation that was performed automatically using Rigaku SAXSLab software. Data were collected in 10-minute frames with each frame processed separately to assess radiation damage; there were no significant changes in the scattering curves over time. Background scattering from water or buffer collected at the same temperature was subtracted from each frame, and the background-corrected $I(q)$ from the individual frames was then averaged, with the standard deviation taken to be the measurement uncertainty and used in weighted least-squares analyses described below.

*Small-angle Neutron Scattering (SANS).* SANS experiments were performed on the NG7 and NGB30 30m SANS instruments at the NIST Center for Neutron Research (NCNR). Data were collected using a neutron wavelength of 6 Å and wavelength spread ($\Delta\lambda/\lambda$) of 0.13. Measurements were made with sample-to-detector distances (SDD) of 1 m, 4 m, and 13 m. Additional low q data were collected with a lenses configuration that used a neutron wavelength of 8.09 Å and SDD of either 13 m or 15 m depending on the instrument. The configurations provided access to an approximate q-range of 0.001 $Å^{-1}$ < q < 0.4 $Å^{-1}$, where q is as described above except λ refers to the neutron wavelength. Data were collected for 1-3 h depending on the vesicle concentration and were corrected for empty cell and instrument background using the IGOR Pro Macros provided by NIST (1).

Vesicles for SANS experiments were prepared at 10 mg/mL or 20 mg/mL as described above. Some samples were also diluted to 2 mg/mL for SANS measurements to minimize the structure factor from inter-vesicle interactions and better determine the vesicle radius and polydispersity at low q.

*Scattering model.* Scattering data were analyzed following (2,3) with modifications described here. The experimentally observed scattering intensity from a vesicle suspension can be expressed as the product of a single-bilayer vesicle form factor $P(q)$ and a structure factor $S(q)$ that accounts for density correlations between different bilayers (e.g., the stacked bilayers in a multilamellar vesicle):

$$I(q) = q^{-2}P(q)S(q)\,. \qquad Eq.S1$$

Pencer et al. (4) showed that, for a polydisperse vesicle suspension whose sizes follow a Schulz distribution, $P(q)$ is well-approximated by

$$P(q) \approx P_V(R_m, \sigma, q)|F_B(q)|^2\,, \qquad Eq.S2$$

where $R_m$ is the average vesicle radius, $\sigma$ is the relative size polydispersity, and

$$P_V(R_m, \sigma, q) = 8\pi^2 {R_m}^2 \sigma^4 (1+\sigma^{-2})(2+\sigma^{-2}) \times \left[1 - \frac{\left(1+4q^2{R_m}^2\sigma^4\right)^{-\frac{1}{2\sigma^2}} \cos\left((2+\sigma^{-2})\tan^{-1}(2qR_m\sigma^2)\right)}{1+4q^2{R_m}^2\sigma^4}\right]. \qquad Eq.S3$$

Equation S2 is valid provided the vesicle radius is much larger than the bilayer thickness. $P_V$ was set to unity in SAXS analyses because vesicle size does not influence SAXS data within the experimental $q$ range of this study.

The flat bilayer scattering amplitude $F_B(q)$ in Eq. S2 accounts for density correlations within a single bilayer, normal to the plane of the bilayer (i.e., the lipid bilayer structure). We model $F_B$ using a symmetric six-slab volume probability distribution with separate components for the lipid headgroups, combined CH and $CH_2$ groups of the hydrocarbon chains, and terminal $CH_3$ groups at the bilayer midplane (2):

$$F_B(q) = \frac{2e^{-\frac{(q\sigma_S)^2}{2}}}{qD_H A_L V_T V_W (V_C - 2V_T)} \Big| V_T\{b_W(A_L D_H - V_H)(V_C - 2V_T) + V_W b_H (V_C - 2V_T) - V_W A_L D_H (b_C - 2b_T)\} \sin\left(\frac{qV_C}{A_L}\right) + V_T(V_C - 2V_T)(b_W V_H - b_H V_W) \sin\left(qD_H + \frac{qV_C}{A_L}\right) + V_W A_L D_H (b_C V_T - b_T V_C) \sin\left(\frac{2qV_T}{A_L}\right) \Big| . \qquad Eq. S4$$

In Eq. S4, $V_C$, $V_H$, $V_T$, and $V_W$ are the molecular volumes of the lipid hydrocarbon chains, headgroup, terminal methyl, and water, respectively, with corresponding scattering factors $b_C$, $b_H$, $b_T$, and $b_W$ (Table S1). To mimic the smoothing effects of thermal disorder, the step-like volume probability profile was convoluted with a Gaussian of width $\sigma_S$ = 2.8 Å. Explicit bilayer structural parameters include the area per lipid $A_L$ and the headgroup thickness $D_H$; additional structural parameters are derived from fitted model parameters, the most important of which are the total (Luzzati) bilayer thickness $D_B = 2(V_C + V_H)/A_L$ and the hydrocarbon thickness $2D_C = 2V_C/A_L$. Because of the large number of model parameters, we constrained $V_T$ (53 Å$^3$) and the width of the headgroup region $D_H$ (6 Å for X-ray data, 10 Å for neutron data unless otherwise stated) to improve the robustness of the fitting routine.

When two or more bilayers are stacked as in an MLV, correlations between the lamellae modulate the scattering according to Bragg's law. The resulting Bragg reflections appear in reciprocal space at multiples of the interlamellar repeat distance *d*, i.e. $q_m = m2\pi/d$ (*m* = 1, 2, 3…), and can be accounted for with a structure factor

$$S_N(q) = N + 2\sum_{k=1}^{N-1}(N-k)\cos(kqd)\, exp[-(dq/2\pi)^2\eta\{\gamma + \ln(\pi k)\}] , \qquad Eq. S5$$

where *N* is the number of bilayers in the stack, $\gamma$ is Euler's constant, and $\eta = \pi k_B T/2d^2\sqrt{Bk_c}$ is the Caillé parameter, which is related to bilayer undulations and depends on the bulk modulus of compression *B* (5) the bilayer bending rigidity $k_c$, and the absolute temperature *T* ($k_B$ is the Boltzmann constant). The width of the Bragg peaks is essentially controlled by $\eta$ and is the result of several types of disorder and crystalline defects that have been thoroughly described elsewhere (3,6).

To account for heterogeneity in the number of stacked lamellae in a vesicle suspension (3), we use a modified exponential distribution in which the relative probability of finding a vesicle with *N* lamellae is given by

$$w_N(\lambda) = \begin{cases} w_1 , & N = 1 \\ e^{-\lambda N}, & N > 1 \end{cases} . \qquad Eq. S6$$

The distribution parameter $\lambda$ is inversely related to the average number of lamellae, and the relative probability $w_1$ of a unilamellar vesicle ($N$ = 1) is a separate parameter; this is a purely empirical modification of the exponential distribution to improve the fit quality. The ensemble-averaged structure factor that is inserted into Eq. S1 is given by

$$\langle S(q) \rangle = \sum_{N=1}^{N_{max}} S_N(q) w_N(\lambda) \;/\; \sum_{N=1}^{N_{max}} w_N(\lambda) \;, \qquad Eq. S7$$

where the (otherwise infinite) sum is truncated at a maximum number of layers governed by a cumulative probability cutoff $p$,

$$N_{max} = \lceil -\log(1-p)/\lambda \rceil \,. \qquad Eq. S8$$

We used a value of $p$ = 0.999 for all analyses presented in this work. The ensemble-averaged number of lamellae $\langle N \rangle$ is obtained from the modified distribution as

$$\langle N \rangle = \sum_{N=1}^{N_{max}} N\, w_N(\lambda) \;/\; \sum_{N=1}^{N_{max}} w_N(\lambda) \;, \qquad Eq. S9$$

and the externally accessible lipid fraction is calculated as

$$f_{acc} = (2\langle N \rangle)^{-1} \,. \qquad Eq. S10$$

With this definition, a perfectly unilamellar sample has $f_{acc}$ = 0.5. Finally, two useful metrics are the number fraction $n_N$ and mass fraction (relative to the total lipid mass) $m_N$ of vesicles with $N$ lamellae:

$$n_N = w_N(\lambda) \;/\; \sum_{N=1}^{N_{max}} w_N(\lambda) \,, \qquad Eq. S11$$

$$m_N = N\, w_N(\lambda) \;/\; \sum_{N=1}^{N_{max}} N\, w_N(\lambda) \,. \qquad Eq. S12$$

*Scattering analysis.* Scattering data were fit using the model described in the previous section, implemented in a nonlinear least-squares routine using custom code written in Mathematica v11.3.0 (Wolfram Research, Champaign, IL). In most cases the adjustable parameters were $A_L$, $V_C$, $\eta$, $d$, $\lambda$, and $w_1$, in addition to a constant background and arbitrary scale factor. We found empirically that the most reliable results were obtained by varying the transformed parameter $\Lambda = \lambda^{-1}$ in a line search and refining the remaining adjustable parameters with a Levenberg-Marquardt algorithm included in Mathematica NonlinearModelFit function.

The robustness of the fitting routine was examined via Monte Carlo error estimation of parameter uncertainties. Briefly, experimental I(q) data were fitted with splines to produce a smooth curve, from which 500 synthetic I(q) datasets were generated by adding Gaussian noise with a standard deviation equal to the $q$-dependent measurement uncertainties determined during data reduction. The synthetic datasets were fit as described above to generate distributions of the best fit parameters, from which standard deviations were calculated. Table S4 shows the parameter mean values $\mu$, standard deviations $\sigma$, and relative errors $\sigma/\mu$ for POPC vesicles hydrated in PBS buffer and extruded with a 100 nm pore size filter. The parameter uncertainty calculated from best fit values obtained from a set of sample replicates is in all cases larger than that calculated for each individual sample from a set of synthetic Monte Carlo replicates, demonstrating that the uncertainty arising from sample-to-sample variation overwhelms the uncertainty inherent to the model fitting.

*Cryo-electron Microscopy (CryoEM).* To cryopreserve vesicles, 4 µL of a 1 mg/mL sample were applied to a Quantifoil 2/2 carbon coated 200 mesh copper grid that was glow-discharged for 1 min at 15 mA in a Pelco Easi-Glow discharge device. After manual blotting, the grids were plunged into liquid ethane cooled with liquid $N_2$. Cryo-preserved grids were stored in liquid $N_2$ until use. Samples were coded prior to preservation and all sample preparation and image collection was accomplished by an investigator blind to the sample composition.

Image collection was accomplished at approximately 2 µm under focus on a FEI Polara G2 operated at 300 kV equipped with a Gatan K2 Summit direct electron detector operated in photon counting mode. Data collection was performed in a semi-automated fashion using Serial EM software operated in low-dose mode (7). Briefly, areas of interest were identified visually and 8x8 montages were collected at low magnification (2400x) at various positions across the grid and then individual areas were marked for automated data collection. Data was collected at 2.51 Å/pixel. Movies of 30 dose-fractionated frames were collected at each target site with the total electron dose being kept to < 20 $e^-/Å^2$. Dose-fractionated movies were drift-corrected with MotionCor2 (8).

*Cryo-electron Microscopy data analysis.* The extent of multillamellarity in LUV preparations was calculated from cryoEM images. For each sample, 120-190 vesicles from at least six images were measured and the number of lamellae manually counted. For preparations that resulted in formation of nested vesicles (i.e. smaller vesicles inside larger vesicles without characteristic interlamellar spacing), diameters of inner nested vesicles were recorded. $f_{acc}$ was calculated in multilamellar vesicles with the assumption that each layer of the MLV had a diameter of 8 nm smaller than the preceding layer. The final calculations were almost insensitive to reasonable variations in this parameter.

### SUPPORTING TABLES

**Table S1** Volumes and scattering factors of lipids used in this study: $V_H$, headgroup volume; $V_C$, hydrocarbon volume; $b_H^X$, headgroup X-ray scattering factor; $b_H^N$, headgroup neutron scattering factor; $b_C^X$, hydrocarbon X-ray scattering factor; $b_C^N$, hydrocarbon neutron scattering factor. X-ray scattering factors are calculated as the total number of electrons of the constituent atoms (neglecting the $q$-dependence), and neutron scattering factors are calculated as the sum of atomic neutron scattering lengths of the constituent atoms.

| Lipid | T [ºC] | $V_H$ [Å$^3$] | $b_H^X$ [e$^-$] | $b_H^N$ [fm Å$^{-3}$] | $V_C$ [Å$^3$] | $b_C^X$ [e$^-$] | $b_C^N$ [fm Å$^{-3}$] |
|---|---|---|---|---|---|---|---|
| DLPC[a] | 25 | 331 | 164 | 60.072 | 656.5 | 178 | -25.782 |
| | 30 | 331 | 164 | 60.072 | 659.8 | 178 | -25.782 |
| DMPC[a] | 30 | 331 | 164 | 60.072 | 768.9 | 210 | -29.11 |
| | 35 | 331 | 164 | 60.072 | 772.8 | 210 | -29.11 |
| DPPC[a] | 50 | 331 | 164 | 60.072 | 897.5 | 242 | -32.438 |
| POPC[a] | 25 | 331 | 164 | 60.072 | 920.5 | 256 | -26.624 |
| | 35 | 331 | 164 | 60.072 | 929.8 | 256 | -26.624 |
| SOPC[a] | 25 | 331 | 164 | 60.072 | 973.5 | 272 | -28.288 |
| DOPC[b] | 30 | 331 | 164 | 60.072 | 972.3 | 270 | -20.81 |
| Di20:1PC[b] | 30 | 331 | 164 | 60.072 | 1079.7 | 302 | -24.138 |
| Di22:1PC[b] | 30 | 331 | 164 | 60.072 | 1189.9 | 334 | -27.466 |
| DMPG/$D_2O$[c] | 35 | 291 | 166 | 95.91 | 766.5 | 210 | -29.11 |
| POPG/$H_2O$[c] | 25 | 291 | 166 | 75.09 | 914.9 | 256 | -26.624 |
| POPS/$H_2O$[d] | 25 | 278 | 172 | 88.189 | 920.5 | 256 | -26.624 |
| Egg-SM/$H_2O$[e] | 50 | 274 | 150 | 47.441 | 883.7 | 240 | -24.96 |

[a]ref. 9; [b]ref. 10; [c]ref. 11; [d]ref. 12; [e]Norbert Kučerka, personal communication.

**Table S2** Vesicle size and polydispersity index (PDI) obtained from DLS, and bilayer structural parameters obtained from SAXS analysis (parameters are defined in Section S1). Values in italics were constrained.

| Composition | Solvent | T [°C] | Extrusion pore size [nm] | *DLS parameters* diam. [nm] | PDI | *SAXS parameters* $\langle N \rangle$ | $f_{acc}$ | $n_1$ | $m_1$ | $d$ [Å] | $\eta$ | $V_L$ [Å$^3$][a] | $A_L$ [Å$^2$] |
|---|---|---|---|---|---|---|---|---|---|---|---|---|---|
| POPC | $H_2O$ | 25 | unextruded | 7600 | 0.13 | 3.42 | 0.15 | 0.29 | 0.09 | 62.6 | 0.09 | 1257 | *65* |
| | | | 1000 | 272 | 0.27 | 1.63 | 0.31 | 0.69 | 0.42 | 63.0 | 0.08 | 1257 | 64.7 |
| | | | 400 | 295 | 0.30 | 1.62 | 0.31 | 0.68 | 0.42 | 63.0 | 0.08 | 1257 | 64.6 |
| | | | 200 | 182 | 0.16 | 1.35 | 0.37 | 0.77 | 0.57 | 63.0 | 0.09 | 1256 | 64.4 |
| | | | 100 | 143 | 0.10 | 1.21 | 0.41 | 0.83 | 0.68 | 63.2 | 0.10 | 1255 | 64.3 |
| | | | | 126 | 0.17 | 1.18 | 0.42 | 0.85 | 0.72 | 63.8 | 0.08 | 1253 | 63.3 |
| | | | 50 | 108 | 0.09 | 1.09 | 0.46 | 0.92 | 0.85 | 63.4 | 0.11 | 1255 | 64.2 |
| | | | 30 | | | 1.03 | 0.48 | 0.97 | 0.94 | 64.5 | 0.07 | 1255 | 63.8 |
| | PBS | 25 | unextruded | | | 6.74 | 0.07 | 0.15 | 0.02 | 63.9 | 0.07 | 1264 | *65* |
| | | | | | | 17.8 | 0.03 | 0.05 | 0.00 | 62.6 | 0.07 | 1252 | *65* |
| | | | | | | 7.81 | 0.06 | 0.13 | 0.02 | 63.8 | 0.08 | 1265 | *65* |
| | | | 100 | 134 | 0.25 | 1.51 | 0.33 | 0.70 | 0.47 | 62.0 | 0.09 | 1262 | 64.8 |
| | | | | 138 | 0.12 | 1.40 | 0.36 | 0.75 | 0.54 | 62.4 | 0.08 | 1263 | 64.6 |
| | | | | 136 | 0.15 | 1.34 | 0.37 | 0.77 | 0.57 | 62.2 | 0.08 | 1262 | 64.3 |
| | | | | 115 | 0.12 | 1.41 | 0.35 | 0.75 | 0.53 | 62.3 | 0.08 | 1262 | 64.4 |
| | $D_2O$ | 35 | 100 | 116 | 0.03 | 1.11 | 0.45 | 0.90 | 0.82 | 63.7 | 0.11 | 1261 | 65.3 |
| POPC/POPG 95/5 | $H_2O$ | 25 | unextruded | 495 | 0.29 | 1.18 | 0.42 | 0.87 | 0.73 | 71.5 | 0.44 | 1257 | 65.0 |
| | | | 1000 | 353 | 0.20 | 1.16 | 0.43 | 0.88 | 0.75 | 73.6 | 0.44 | 1256 | 64.7 |
| | | | 400 | 261 | 0.15 | 1.16 | 0.43 | 0.88 | 0.76 | 74.4 | 0.53 | 1255 | 64.4 |
| | | | 200 | 172 | 0.10 | 1.07 | 0.47 | 0.93 | 0.87 | 75.3 | 0.52 | 1253 | 63.9 |
| | | | 100 | 135 | 0.10 | 1.03 | 0.48 | 0.97 | 0.93 | *75* | *0.52* | 1253 | 63.8 |
| | | | 50 | 103 | 0.09 | 1.02 | 0.49 | 0.99 | 0.97 | *75* | *0.52* | 1252 | 63.7 |
| | | | 30 | 72 | 0.08 | 1.01 | 0.49 | 0.99 | 0.98 | *75* | *0.52* | 1253 | 63.8 |
| POPC/POPG 98/2 | $D_2O$ | 35 | 100 | 130 | 0.04 | 1.03 | 0.49 | 0.97 | 0.95 | 71.6 | *0.40* | 1262 | 65.5 |
| POPC/POPG 96/4 | | | | 125 | 0.11 | 1.02 | 0.49 | 0.98 | 0.96 | 76.8 | *0.50* | 1262 | 65.5 |
| POPC/POPG 94/6 | | | | 120 | 0.10 | 1.02 | 0.49 | 0.98 | 0.95 | *75* | *0.50* | 1261 | 65.4 |
| POPC/POPG 92/8 | | | | 130 | 0.09 | 1.03 | 0.48 | 0.98 | 0.95 | *75* | *0.50* | 1261 | 65.3 |
| POPC/POPG 90/10 | | | | 117 | 0.12 | 1.03 | 0.49 | 0.97 | 0.95 | *75* | *0.50* | 1261 | 65.4 |
| POPC/POPG 98/2 | PBS | 25 | 100 | 127 | 0.21 | 1.36 | 0.37 | 0.76 | 0.56 | 63.3 | 0.11 | 1264 | 64.7 |
| POPC/POPG 96/4 | | | | 113 | 0.21 | 1.18 | 0.42 | 0.85 | 0.72 | 66.6 | 0.17 | 1262 | 64.5 |
| POPC/POPG 94/6 | | | | 118 | 0.24 | 1.14 | 0.44 | 0.88 | 0.77 | 70.7 | 0.25 | 1262 | 64.5 |
| POPC/POPG 92/8 | | | | 94 | 0.27 | 1.11 | 0.45 | 0.91 | 0.82 | 71.8 | 0.25 | 1262 | 64.4 |
| POPC/POPG 90/10 | | | | 100 | 0.28 | 1.10 | 0.46 | 0.91 | 0.83 | *75* | 0.38 | 1261 | 64.5 |
| POPC/POPS 98/2 | $H_2O$ | 25 | 100 | 115 | 0.24 | 1.01 | 0.50 | 0.85 | 0.98 | *63* | *0.2* | 1253 | 63.3 |
| POPC/POPS 96/4 | | | | 108 | 0.21 | 1.00 | 0.50 | 0.99 | 1.0 | *63* | *0.2* | 1252 | 63.1 |
| POPC/POPS 94/6 | | | | 115 | 0.14 | 1.00 | 0.50 | 1.0 | 1.0 | *63* | *0.2* | 1252 | 63.0 |
| POPC/POPS 92/8 | | | | 108 | 0.31 | 1.00 | 0.50 | 1.0 | 1.0 | *63* | *0.2* | 1250 | 62.9 |
| POPC/POPS 90/10 | | | | 118 | 0.24 | 1.00 | 0.50 | 1.0 | 1.0 | *63* | *0.2* | 1249 | 62.7 |
| DLPC | $H_2O$ | 30 | unextruded | 1600 | 0.15 | 12.7 | 0.04 | 0.08 | 0.01 | 57.7 | 0.10 | 990 | *62.1* |
| | | | 100 | 113 | 0.08 | 1.14 | 0.44 | 0.88 | 0.77 | 58.6 | 0.13 | 994 | 62.1 |
| DMPC | | 30 | unextruded | 1800 | 0.23 | 10.2 | 0.05 | 0.10 | 0.01 | 62.4 | 0.09 | 1100 | *58* |
| | | | 100 | 102 | 0.06 | 1.10 | 0.45 | 0.91 | 0.82 | 63.7 | 0.15 | 1099 | 58.0 |
| DPPC | | 50 | unextruded | 2300 | 0.005 | 5.9 | 0.08 | 0.16 | 0.03 | 65.3 | 0.10 | 1237 | *62.4* |
| | | | 100 | 172 | 0.18 | 1.09 | 0.46 | 0.92 | 0.84 | 67.8 | 0.16 | 1227 | 62.4 |
| Egg-SM | | 50 | unextruded | 522 | 0.36 | 1.18 | 0.42 | 0.98 | 0.84 | 62.5 | 0.05 | 1178 | 61.0 |
| | | | 100 | 114 | 0.04 | 1.02 | 0.49 | 0.99 | 0.97 | 66.1 | 0.02 | 1177 | 60.6 |
| SOPC | | 25 | unextruded | | | 7.5 | 0.07 | 0.13 | 0.02 | 65.2 | 0.05 | 1302 | 61.8 |
| | | | 100 | 127 | 0.03 | 1.20 | 0.42 | 0.84 | 0.70 | 65.8 | 0.08 | 1308 | 63.1 |
| | | | | 128 | 0.19 | 1.27 | 0.39 | 0.80 | 0.62 | 65.6 | 0.08 | 1309 | 63.3 |
| DOPC | | 30 | unextruded | 1500 | 0.005 | 9.4 | 0.05 | 0.10 | 0.01 | 62.4 | 0.07 | 1294 | *68.9* |
| | | | 100 | 130 | 0.12 | 1.16 | 0.43 | 0.86 | 0.74 | 63.4 | 0.10 | 1302 | 68.9 |
| Di20:1-PC | | 30 | unextruded | 2620 | 0.17 | 5.2 | 0.10 | 0.19 | 0.04 | 65.4 | 0.06 | 1412 | *67.1* |
| | | | 100 | 120 | 0.05 | 1.08 | 0.46 | 0.93 | 0.86 | 67.2 | 0.06 | 1412 | 67.1 |
| Di22:1-PC | | 30 | unextruded | 2000 | 0.005 | 9.2 | 0.05 | 0.11 | 0.01 | 69.3 | 0.03 | 1515 | *64.8* |
| | | | 100 | 140 | 0.02 | 1.21 | 0.41 | 0.83 | 0.68 | 70.3 | 0.04 | 1520 | 64.8 |

[a]The lipid hydrocarbon volume $V_C$ was allowed to vary from the initial input values given in Table S1; we report the best-fit total lipid volume $V_L = V_C + V_H$ here.

**Table S3** Vesicle size distribution and bilayer structural parameters obtained from SANS analysis (parameters are defined in Section S1). Values in italics were constrained.

| Composition | Solv. | T [°C] | Extrusion pore size [nm] | Conc. [mg/mL] | *SANS parameters* | | | | | | | | |
|---|---|---|---|---|---|---|---|---|---|---|---|---|---|
| | | | | | diam. [nm] | $\sigma$ | $\langle N \rangle$ | $f_{acc}$ | $n_1$ | $m_1$ | $d$ [Å] | $\eta$ | $A_L$ [Å$^2$] |
| DLPC | $D_2O$ | 25 | 400 | 20 | 342 | 0.13 | 2.45 | 0.20 | 0.54 | 0.22 | 58.1 | 0.18 | *61* |
| | | | | 2 | 614 | 0.17 | 3.00 | 0.17 | 0.44 | 0.15 | 57.5 | 0.22 | *61* |
| | | | 200 | 20 | 205 | 0.26 | 1.44 | 0.35 | 0.77 | 0.54 | 59.1 | 0.19 | *61* |
| | | | | 2 | 131 | 0.30 | 1.64 | 0.30 | 0.67 | 0.41 | 57.0 | 0.28 | *61* |
| | | | 100 | 20 | 71 | 0.42 | 1.20 | 0.42 | 0.85 | 0.71 | 58.0 | 0.40 | 61.0 |
| | | | | 2 | 87 | 0.26 | 1.25 | 0.40 | 0.83 | 0.67 | 57.8 | 0.33 | 59.9 |
| | | | 50 | 20 | 51 | 0.31 | 1.03 | 0.48 | 0.97 | 0.94 | 63.4 | 0.82 | 62.8 |
| | | | | 2 | 53 | 0.26 | 1.06 | 0.47 | 0.95 | 0.89 | *62* | *0.8* | 60.5 |
| | | | 30 | 20 | 49 | 0.33 | 1.05 | 0.48 | 0.96 | 0.91 | 63.0 | 0.64 | 61.3 |
| | | | | 2 | 53 | 0.28 | 1.09 | 0.46 | 0.94 | 0.86 | 61.0 | 0.62 | *61* |
| DMPC | $D_2O$ | 35 | 400 | 20 | 711 | 0.16 | 2.35 | 0.21 | 0.60 | 0.26 | 61.8 | 0.24 | 59.7 |
| | | | 200 | | 131 | 0.35 | 1.47 | 0.34 | 0.78 | 0.53 | 62.5 | 0.23 | 59.5 |
| | | | 100 | | 96 | 0.32 | 1.20 | 0.42 | 0.91 | 0.76 | 62.1 | 0.11 | *59* |
| | | | 50 | | 59 | 0.33 | 1.09 | 0.46 | 0.96 | 0.88 | 67.1 | 0.56 | *59* |
| | | | 30 | | 52 | 0.33 | 1.08 | 0.46 | 0.96 | 0.89 | *63* | 0.52 | *59* |
| DMPC | $D_2O$/ 100 mM NaCl | 35 | 100 | 10 | 92 | 0.30 | 1.19 | 0.42 | 0.84 | 0.71 | 62.4 | 0.45 | 59.1 |
| DMPC/DMPG 98/2 | | | | | 92 | 0.33 | 1.40 | 0.36 | 0.80 | 0.57 | 68.1 | 0.57 | 59.3 |
| DMPC/DMPG 95/5 | | | | | 91 | 0.30 | 1.14 | 0.44 | 0.90 | 0.79 | 98.5 | 1.17 | 58.7 |
| DMPC/DMPC 90/10 | | | | | 86 | 0.29 | 1.16 | 0.43 | 0.87 | 0.76 | 113.9 | 0.75 | 58.3 |
| DMPC/DMPG 80/20 | | | | | 81 | 0.33 | 1.13 | 0.44 | 0.89 | 0.79 | 126.8 | 0.72 | 58.7 |
| DMPC/DMPG 50/50 | | | | | 108 | 0.32 | 1.13 | 0.44 | 0.89 | 0.80 | 138.3 | 0.71 | 59.8 |
| DMPG | | | | | 87 | 0.34 | 1.06 | 0.47 | 0.94 | 0.89 | 156.3 | 0.43 | 62.6 |

**Table S4** Uncertainty in fitted model parameters for SAXS data from POPC hydrated in PBS buffer and extruded with a 100 nm pore size filter (parameters are defined in Section S1). Columns 2-4 correspond to the average $\mu$, standard deviation $\sigma$, and percent error (defined here as 100 $\sigma/\mu$) from four replicate samples. The remaining columns show the same information calculated for each of the individual samples using Monte Carlo error estimation as described in the Materials and Methods.

| | Sample replicates (N = 4) | | | Sample 1 MC replicates (N = 500) | | | Sample 2 MC replicates (N = 500) | | | Sample 3 MC replicates (N = 500) | | | Sample 4 MC replicates (N = 500) | | |
|---|---|---|---|---|---|---|---|---|---|---|---|---|---|---|---|
| Param | $\mu$ | $\sigma$ | % err | $\mu$ | $\sigma$ | % err | $\mu$ | $\sigma$ | % err | $\mu$ | $\sigma$ | % err | $\mu$ | $\sigma$ | % err |
| $A_L$ | 64.5 | 0.2 | **0.3** | 64.8 | 0.10 | **0.15** | 64.6 | 0.05 | **0.08** | 64.4 | 0.05 | **0.08** | 64.5 | 0.05 | **0.08** |
| $V_C$ | 931.2 | 0.5 | **0.05** | 930.7 | 0.3 | **0.03** | 931.8 | 0.15 | **0.02** | 931.1 | 0.13 | **0.01** | 931.7 | 0.14 | **0.02** |
| $\eta$ | 0.082 | 0.008 | **9.5** | 0.093 | 0.003 | **2.8** | 0.082 | 0.002 | **2.0** | 0.077 | 0.002 | **2.2** | 0.079 | 0.002 | **2.1** |
| $d$ | 62.2 | 0.2 | **0.3** | 61.98 | 0.015 | **0.02** | 62.40 | 0.01 | **0.02** | 62.20 | 0.01 | **0.02** | 62.27 | 0.01 | **0.02** |
| $\langle N \rangle$ | 1.42 | 0.07 | **4.9** | 1.51 | 0.006 | **0.4** | 1.40 | 0.003 | **0.2** | 1.34 | 0.002 | **0.2** | 1.41 | 0.003 | **0.2** |
| $f_{acc}$ | 0.35 | 0.02 | **4.8** | 0.33 | 0.001 | **0.4** | 0.36 | 0.001 | **0.2** | 0.37 | 0.001 | **0.2** | 0.35 | 0.001 | **0.2** |
| $n_1$ | 0.74 | 0.03 | **3.7** | 0.70 | 0.004 | **0.5** | 0.74 | 0.002 | **0.5** | 0.77 | 0.002 | **0.2** | 0.74 | 0.002 | **0.3** |
| $m_1$ | 0.53 | 0.04 | **8.4** | 0.46 | 0.004 | **0.9** | 0.53 | 0.002 | **0.5** | 0.57 | 0.002 | **0.4** | 0.53 | 0.002 | **0.5** |

**Table S5** Lipid and membrane physical properties and correlation coefficients with the externally accessible lipid fraction $f_{acc}$ for extruded 100 nm vesicles in pure water.

| Lipid | $f_{acc}$ | Number of carbons per chain | Number of double bonds | Area per lipid [Å$^2$] | Hydrocarbon thickness [Å] | Melting temp. [°C] | Bending modulus [$k_BT$][c] |
|---|---|---|---|---|---|---|---|
| DLPC | 0.44 | 12 | 0 | 62.1 | 21.3 | 7.0 | 25.8 |
| DMPC | 0.45 | 14 | 0 | 58.0 | 26.5 | 23.9 | 34.7 |
| DPPC | 0.46 | 16 | 0 | 62.4 | 28.7 | 41.4 | 34.1 |
| ESM | 0.49 | 17 | 1 | 60.6 | 29.8 | 39.0 | 41.8 |
| POPC | 0.42 | 17 | 1 | 63.3 | 29.1 | –2.0 | 24.3 |
| SOPC | 0.40 | 18 | 1 | 63.3 | 30.9 | 6.0 | 26.4 |
| **r[a]** | | **–0.13** | **–0.18** | **–0.56** | **–0.03** | **0.84** | **0.91** |
| DOPC | 0.43 | 18 | 2 | 68.9 | 28.2 | –17.3 | 18.3 |
| di20:1-PC | 0.46 | 20 | 2 | 67.1 | 32.2 | –4.3 | 21.1 |
| di22:1-PC | 0.41 | 22 | 2 | 64.8 | 36.7 | 13.2 | 26.9 |
| **r[b]** | | **–0.24** | **–0.22** | **–0.31** | **–0.20** | **0.55** | **0.61** |

[a]correlation coefficient considering only mixed-chain and fully saturated lipids; [b]correlation coefficient considering all lipids; [c]obtained from atomistic simulations in (13) except for diC20:1-PC and diC22:1-PC, which were from a personal communication (M. Doktorova).

### SUPPORTING FIGURES

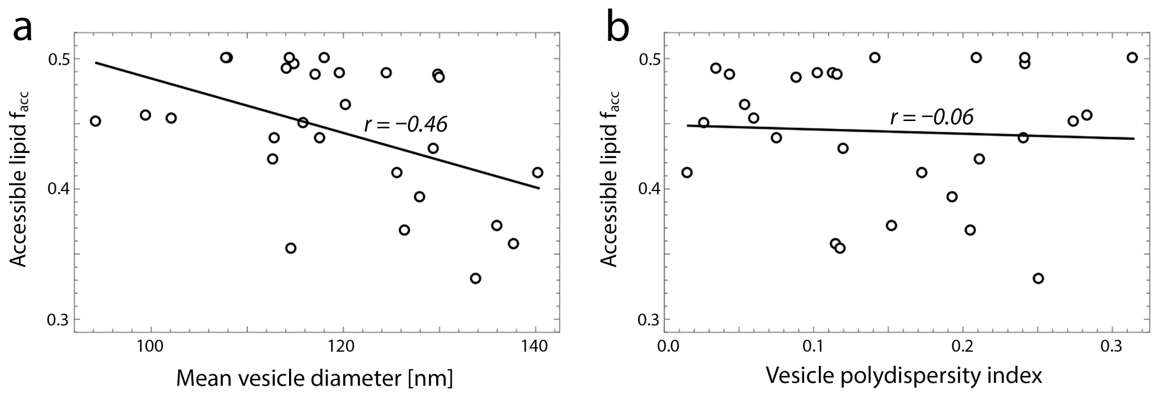

**Figure S1 DLS is insensitive to multilamellar vesicles in extruded samples.** For vesicles extruded with a 100 nm pore size filter (Table S2), the fraction of externally accessible lipid $f_{acc}$ determined from SAXS is weakly correlated with the mean vesicle diameter (Pearson correlation coefficient *r* = –0.46) and is uncorrelated with the vesicle polydispersity index (*r* = –0.06) measured with dynamic light scattering, as shown in panels *a* and *b*, respectively.

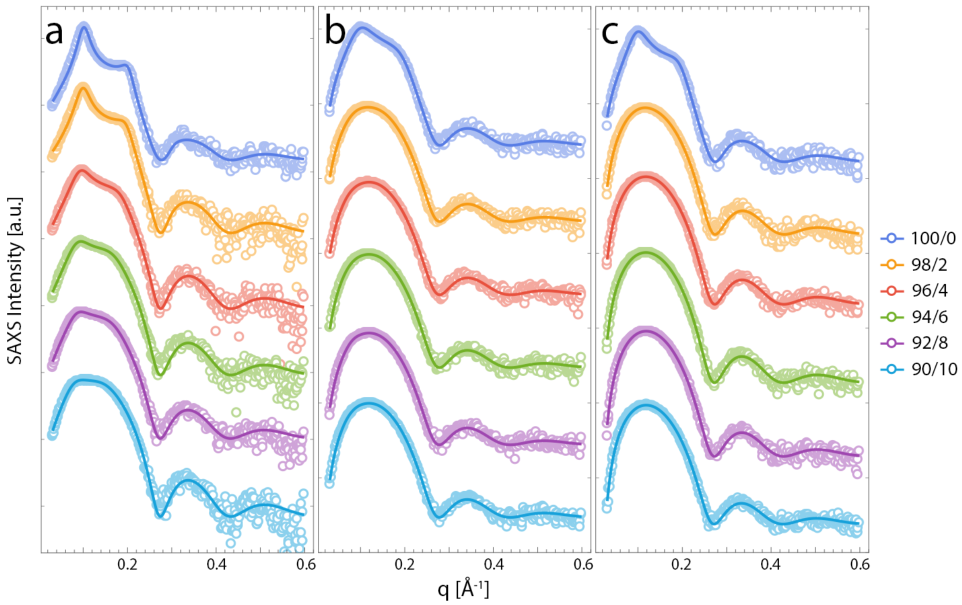

**Figure S2 SAXS data from extruded POPC vesicles with increasing amounts of charged lipid.** (a), POPC/POPG mixtures (molar ratios indicated in the legend) dispersed in PBS buffer at 20 mg/mL and measured at 25°C. (b), POPC/POPG mixtures (molar ratios indicated in the legend) dispersed in $D_2O$ at 20 mg/mL and measured at 35°C. (c), POPC/POPS mixtures (molar ratios indicated in the legend) dispersed in $H_2O$ at 20 mg/mL and measured at 25°C. All vesicles were extruded with a 100 nm pore size filter. Experimental data are shown as open circles and the fitted model is shown as a solid line. Values for fitted parameters are given in Table S2.

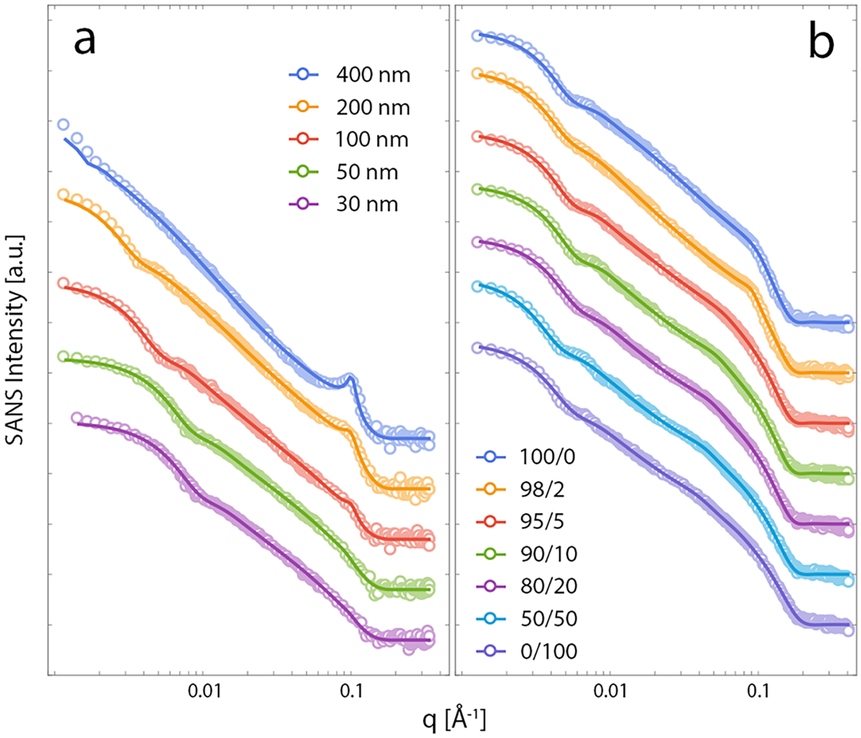

**Figure S3 SANS data from DMPC vesicles.** (a), DMPC dispersed in $D_2O$ at 20 mg/mL, extruded with pore size indicated in legend, and measured at 35°C (b), DMPC/DMPG mixtures (molar ratios indicated in the legend) dispersed in $D_2O$/100 mM NaCl at 10 mg/mL, extruded with a 100 nm pore size filter, and measured at 35°C. Experimental data are shown as open circles and the fitted model is shown as a solid line. Values for fitted parameters are given in Table S3.

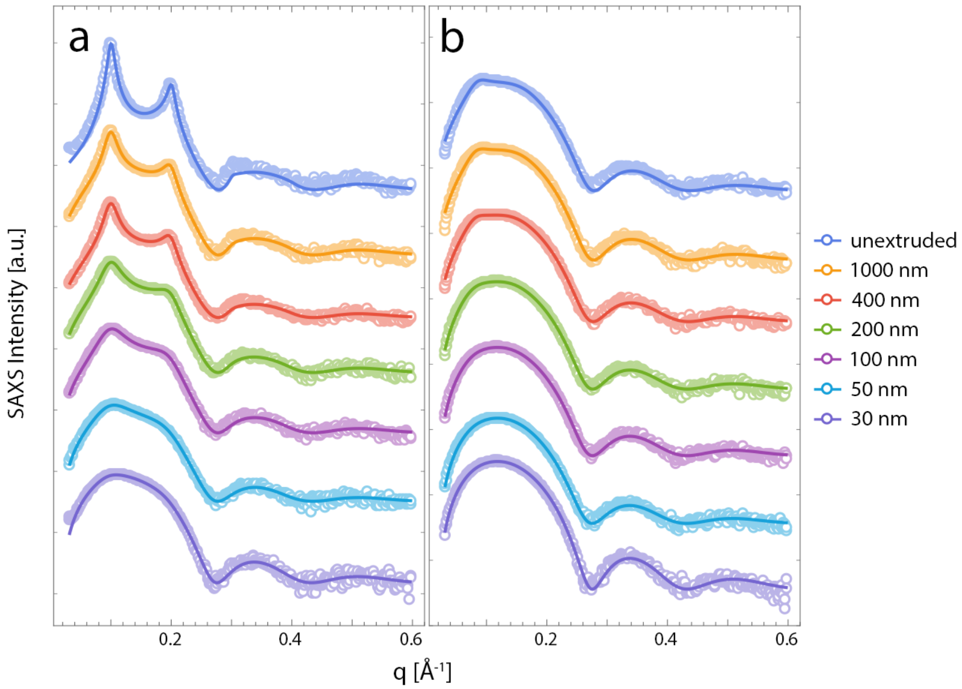

**Figure S4 SAXS data from POPC vesicles extruded through different pore sizes.** POPC (panel *a*) or POPC/POPG 95/5 molar mixture (panel *b*) dispersed in $H_2O$ at 20 mg/mL, extruded with pore size indicated in legend, and measured at 25°C. Experimental data are shown as open circles and the fitted model is shown as a solid line. Values for fitted parameters are given in Table S2.

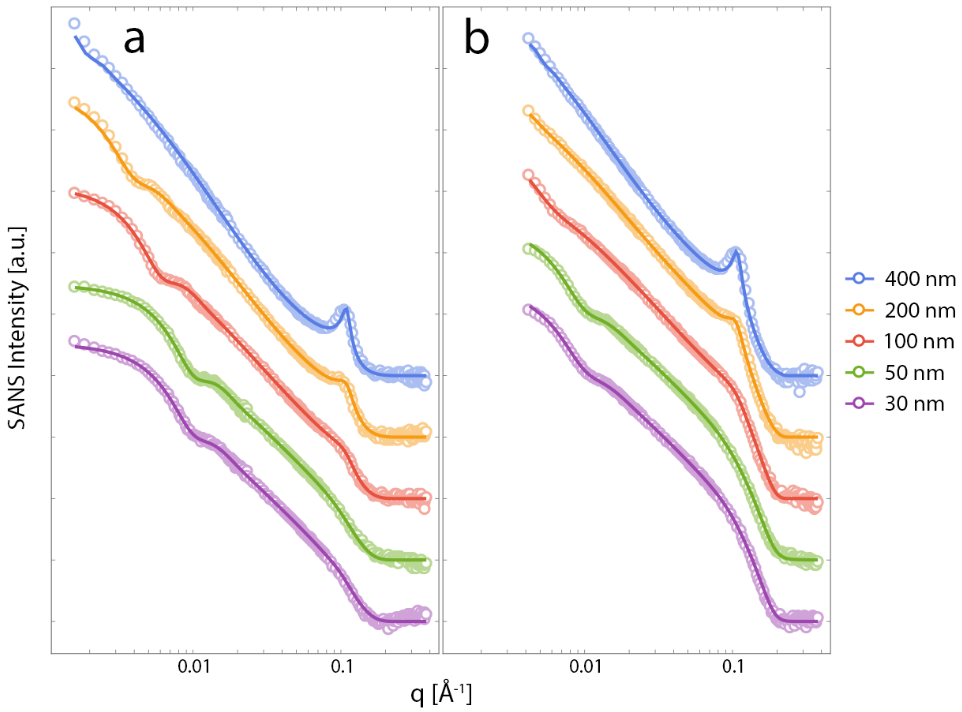

**Figure S5 SANS data from DLPC vesicles extruded through different pore sizes.** DLPC at 2 mg/mL (panel *a*) or 20 mg/mL (panel *b*) dispersed in $D_2O$, extruded with pore size indicated in legend, and measured at 25°C. Experimental data are shown as open circles and the fitted model is shown as a solid line. Values for fitted parameters are given in Table S3.

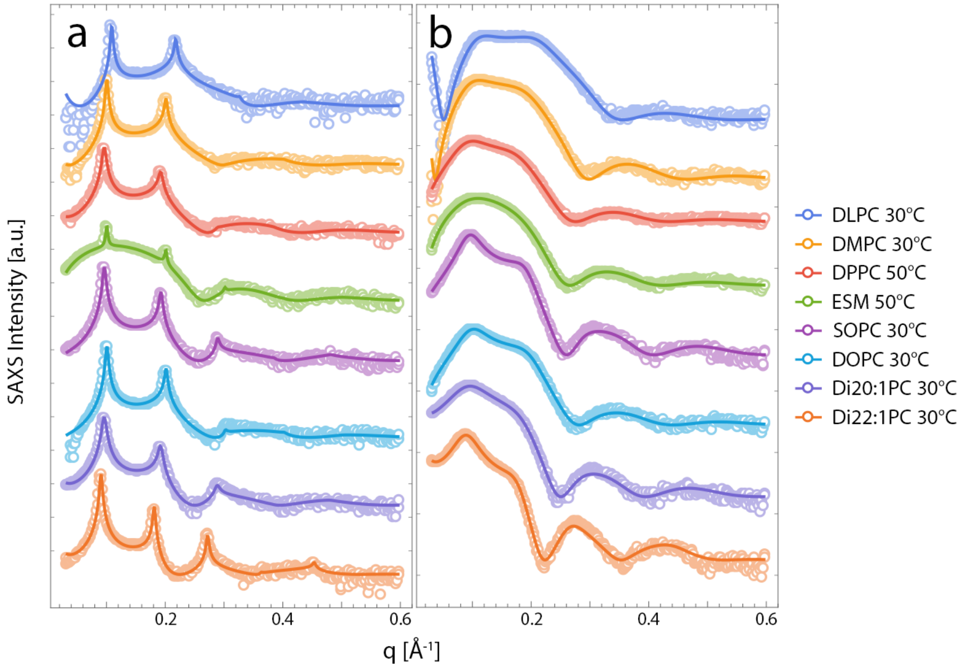

**Figure S6 SAXS data from extruded vesicles composed of neutral lipids in the fluid phase.** Lipid dispersed in $H_2O$ at 20 mg/mL and measured at the temperature indicated in the legend either prior to extrusion (panel *a*) or after extrusion with a 100 nm pore size filter (panel *b*). Experimental data are shown as open circles and the fitted model is shown as a solid line. Values for fitted parameters are given in Table S2.

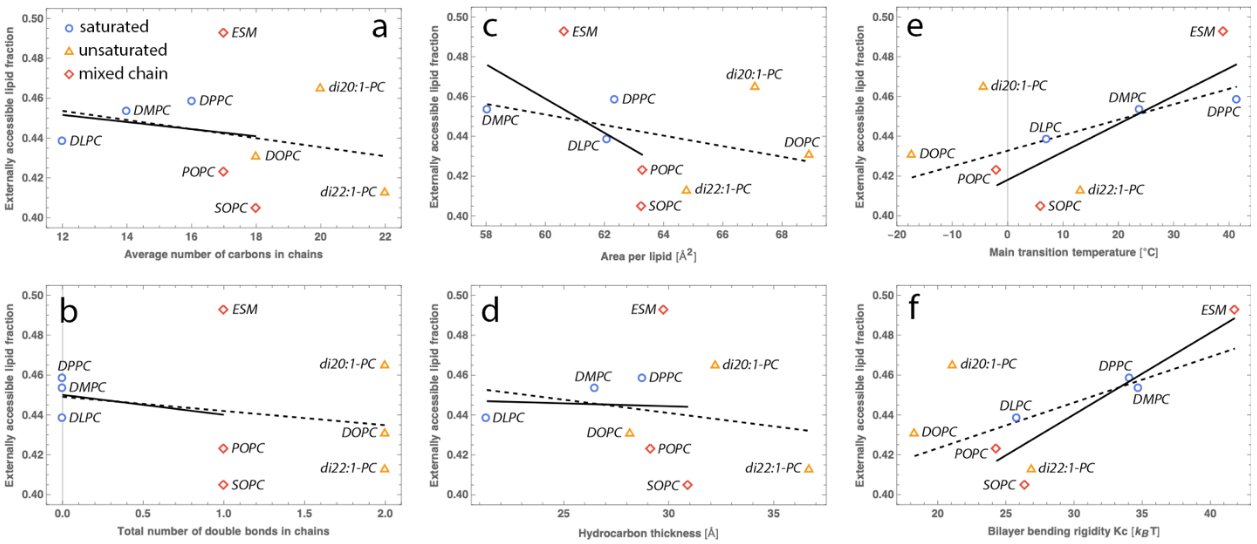

**Figure S7 Average vesicle lamellarity is correlated with membrane bending rigidity.** Shown for various neutral lipids hydrated in water and extruded through 100 nm pores (as indicated in the plots) is the trend between $f_{acc}$ and different membrane physical parameters: (a) the average number of carbons in the chains; (b) the total number of double bonds in the chains; (c) the area per lipid (Table S2); (d) the bilayer's hydrocarbon thickness ($2D_C = 2V_C/A_L$, Table S2); (e) the main transition temperature; and (f) the membrane bending rigidity determined from atomistic molecular simulations. The dashed and solid lines are linear trendlines with and without the di-monounsaturated lipids (i.e., DOPC, di20:1-PC, and di22:1-PC), respectively; in both cases, the strongest correlation is with the bending rigidity. Parameter values and correlation coefficients are given in Table S5.

### REFERENCES

1. S. R. Kline. 2006. Reduction and analysis of SANS and USANS data using IGOR Pro. J. Appl. Crystallogr. 39:895-900.
2. Doktorova, M., Heberle, F. A., Marquardt, D., Rusinova, R., Sanford, R. L., Peyear, T. A., Katsaras, J., Feigenson, G. W., Weinstein, H., and O. S. Andersen. 2019. Gramicidin Increases Lipid Flip-Flop in Symmetric and Asymmetric Lipid Vesicles. Biophys. J. 116:860-873.
3. Pabst, G., Koschuch, R., Pozo-Navas, B., Rappolt, M., Lohner, K., and P. Laggner. 2003. Structural analysis of weakly ordered membrane stacks. J. Appl. Crystallogr. 36:1378-1388.
4. Pencer, J., Krueger, S., Adams, C. P., and J. Katsaras. 2006. Method of separated form factors for polydisperse vesicles. J. Appl. Crystallogr. 39:293-303.
5. De Gennes, P. G., and J. Prost. 1993. The Physics of Liquid Crystals. 2nd ed. Oxford University Press, Oxford, UK.
6. Pabst, G., Rappolt, M., Amenitsch, H., and P. Laggner. 2000. Structural information from multilamellar liposomes at full hydration: full q-range fitting with high quality x-ray data. Phys. Rev. E 62:4000-4009.
7. D. N. Mastronarde. 2005. Automated electron microscope tomography using robust prediction of specimen movements. J. Struct. Biol. 152:36-51.

8. Zheng, S. Q., Palovcak, E., Armache, J. P., Verba, K. A., Cheng, Y., and D. A. Agard. 2017. MotionCor2: anisotropic correction of beam-induced motion for improved cryo-electron microscopy. Nat. Methods 14:331-332.
9. Kučerka, N., Nieh, M. P., and J. Katsaras. 2011. Fluid phase lipid areas and bilayer thicknesses of commonly used phosphatidylcholines as a function of temperature. Biochim. Biophys. Acta 1808:2761-2771.
10. Kučerka, N., Gallova, J., Uhrikova, D., Balgavy, P., Bulacu, M., Marrink, S. J., and J. Katsaras. 2009. Areas of monounsaturated diacylphosphatidylcholines. Biophys. J. 97:1926-1932.
11. Pan, J., Heberle, F. A., Tristram-Nagle, S., Szymanski, M., Koepfinger, M., Katsaras, J., and N. Kučerka. 2012. Molecular structures of fluid phase phosphatidylglycerol bilayers as determined by small angle neutron and X-ray scattering. Biochim. Biophys. Acta 1818:2135-48.
12. Pan, J., Cheng, X., Monticelli, L., Heberle, F. A., Kučerka, N., Tieleman, D. P., and J. Katsaras. 2014. The molecular structure of a phosphatidylserine bilayer determined by scattering and molecular dynamics simulations. Soft Matter 10:3716-3725.
13. Doktorova, M., Harries, D., and G. Khelashvili. 2017. Determination of bending rigidity and tilt modulus of lipid membranes from real-space fluctuation analysis of molecular dynamics simulations. Phys. Chem. Chem. Phys. 19:16806-16818.
